# Cosmos: A Position-Resolution Causal Model for Direct and Indirect Effects in Protein Functions

**DOI:** 10.1101/2025.08.01.667517

**Authors:** Jingyou Rao, Mingsen Wang, Matthew Howard, Willow Coyote-Maestas, Harold Pimentel

**Affiliations:** Department of Computer Science, UCLA, Los Angeles, CA, USA; Department of Mathematics, Baruch College, CUNY, New York City, NY, USA; Department of Bioengineering and Therapeutic Sciences, UCSF, San Francisco, CA, USA; Tetrad Graduate Program, UCSF, San Francisco, CA, USA; Quantitative Biosciences Institute, UCSF, San Francisco, CA, USA; Department of Computational Medicine, David Geffen School of Medicine, UCLA, Los Angeles, CA, USA; Department of Human Genetics, David Geffen School of Medicine, UCLA, Los Angeles, CA, USA

**Keywords:** Bayesian Model Selection, Causal Inference, Deep Mutational Scanning, Multi-Phenotype, Protein Functions

## Abstract

Multi-phenotype deep mutational scanning (DMS) experiments provide a powerful means to dissect how protein variants affect different layers of molecular function, such as abundance, surface expression, and ligand binding. When these phenotypes are connected through a molecular pathway, interpreting variant effects becomes challenging because downstream phenotypes often reflect both direct and indirect consequences of mutation. We introduce `Cosmos`, a Bayesian framework for residue-level causal inference in multi-phenotype DMS data. `Cosmos` addresses three key questions: (1) whether a causal relationship exists between two phenotypes; (2) the strength of that relationship; and (3) the expected downstream phenotype if the upstream phenotype were normalized, enabling counterfactual interpretation. The framework uses position-level aggregation and Bayesian model selection to infer interpretable causal structures, without requiring phenotype-specific biophysical assumptions. We apply `Cosmos` to three datasets—Kir2.1 (abundance and surface expression), PSD95-PDZ3 (abundance and CRIPT binding), and KRAS (abundance and RAF1-RBD binding) and show that it effectively distinguishes direct from indirect functional effects. Across these applications, `Cosmos` provides a generalizable and interpretable approach to disentangle causal relationships in high-throughput protein functional screens.

## 1 Introduction

Recent advancements in deep mutational scanning (DMS) have made it increasingly feasible to perform multiple functional screens on the same protein [1,3,7,9,16,17,20]. These multi-phenotype experiments generate richer datasets and offer deeper insights into how protein mutations affect different protein functions (Figure 1A). One common experimental design captures distinct phenotypes along a molecular pathway, such as protein abundance, surface expression, and membrane activity, each reflecting a different functional layer [10,2]. In such a design, phenotypes are often positioned upstream or downstream of one another within a directed molecular cascade. This study aims to develop a computational method to determine whether a mutation impacts a protein function directly, or indirectly through effects on upstream phenotypes.

**Fig 1:**
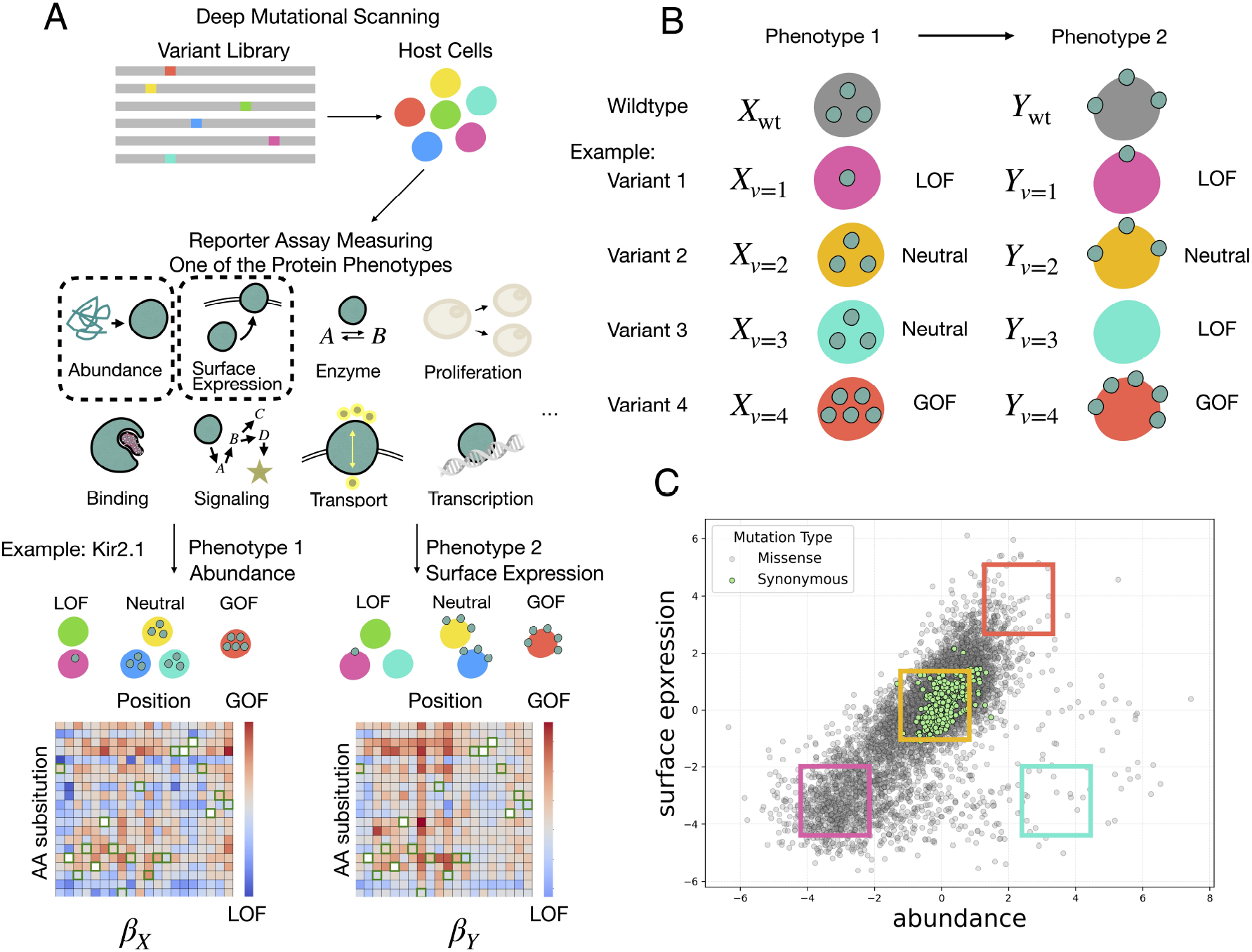
Introduction to multi-phenotype DMS. A: DMS procedure: Variant library is introduced into host cells and various protein phenotypes can be measured by reporter assays. For example, the abundance and surface expression of Kir 2.1 can be measured. The measurements are analyzed and compiled into heatmaps - blue and red representing LOF and GOF respectively. B: The phenotypes of a variant could be in similar or different directions. C: A scatterplot of two phenotypes of the same protein. Colored regions correspond to variant types in B.

Currently, two main approaches are commonly used to analyze such multi-phenotype DMS data. The first is an empirical approach, which fits a regression line, linear [7] or nonlinear [20], using either all variants or a subset such as synonymous mutations, and then calculates residuals to identify variants that deviate from the expected trend. A major challenge in this method is defining how much deviation is meaningful; selecting a threshold is inherently subjective, as we often lack a clear understanding of the expected proportion of true outliers. The second approach employs structured modeling frameworks, such as Mochi [5], which uses biophysical equations to model interactions between latent variables across two phenotypes. While this method offers mechanistic insight, it is highly phenotype-specific, often limited to well-characterized processes like protein folding and binding, and depends heavily on prior knowledge of the phenotypes involved.

Given the sequential nature of these phenotypes, causal analysis becomes a natural approach. Inspired by the Mendelian Randomization framework CAUSE [11], we developed `Cosmos`, a Bayesian model selection framework designed to support causal inference between related phenotypes in deep mutational scanning data with single mutations. `Cosmos` serves three primary purposes: (1) to determine whether a relationship exists between two phenotypes; (2) to estimate the strength of that relationship, i.e., how changes in the upstream phenotype affect the downstream phenotype; and (3) to generate counterfactual predictions—what would happen to the downstream phenotype if the upstream phenotype were fixed to a reference value. Together, these capabilities distinguish direct and indirect effects between related protein functions and offer insights into the causal mechanisms. We first demonstrate `Cosmos` using the Kir2.1 protein, analyzing abundance and surface expression as a vignette to showcase the model’s utility. We then apply the framework to two additional datasets - PSD95-PDZ abundance and CRIPT binding [4], KRAS abundance and RAF1-RBD binding [21] - to further validate its generalizability.

## 2 Results

### 2.1 Multi-Phenotype DMS Experimental Workflow

In a multi-phenotype deep mutational scanning (DMS) experiment, a DMS variant library is first generated. The library often consists of the full spectrum of protein-coding variants that perturb the genotype, including synonymous, missense, and nonsense mutations, as well as insertions, deletions [10], and higher-order combinatorial mutations [12]. This library is then introduced into a biological system suitable for the assay, such as human cells, yeasts, or viruses.

Protein function is typically assessed using a reporter assay that provides an indirect readout, often through proxies like cell proliferation, fluorescence, or other phenotype-specific signals. A wide range of molecular phenotypes can be measured using this approach, including protein abundance [4], surface expression [7,10,22], enzymatic activity [18], cell proliferation [3], ligand binding [4], cell signaling [7], drug or ion transport [2,22], and transcriptional regulation [15] (Fig. 1A). The choice of phenotypes depends on the protein of interest and the specific biological question being addressed.

For example, consider Kir2.1, a potassium channel protein [2,10]. Mutations in Kir2.1 can disrupt protein abundance through effects on folding, stability, or degradation, or impair surface expression due to altered trafficking, membrane insertion, or tetramer complex assembly. These phenotypes can be quantified in two independent reporter assay screens using the same variant library, producing phenotype-specific functional scores for each mutation with statistical methods such as Enrich2 [14], DiMSum [6], `Rosace` [13], and Lilace (Fig. 1A).

Visualizing the data as a heatmap (Fig. 1A) or a scatterplot (Fig. 1C) enables classification of variant effects across phenotypes. Individual variants may exhibit consistent effects across both phenotypes (e.g., variant 1 is loss-of-function (LOF) in both phenotypes—pink; variant 2 is neutral in both—yellow; variant 4 is gain-of-function (GOF) in both, Fig. 1B), or they may differ (e.g., variant 3 is neutral in phenotype 1 but LOF in phenotype 2—cyan, Fig. 1B). These variants will occupy different regions in a phenotype–phenotype scatterplot, helping to reveal the functional relationships between protein phenotypes (Fig. 1C).

### 2.2 Position Aggregation in Cosmos Framework

To infer a causal model under an idealized scenario, we might expect (1) data which cleanly isolates upstream and downstream effects or (2) if not separated, provides multiple measurements that enable inference of the relationships at individual variant *v* resolution. However, given current experimental and practical limitation, further assumptions must be made to infer a tractable causal model from current data.

In situation (1), while upstream phenotypes, such as protein abundance, can often be directly measured, isolating downstream phenotypes without confounding from upstream variation is substantially more difficult. This is because downstream measurements, such as surface expression, reflect a combination of both direct and indirect influences. For example, a mutation might directly impair membrane trafficking (a direct effect on surface expression), or it might reduce surface levels indirectly by destabilizing protein folding, thereby lowering overall protein abundance and limiting the pool available for membrane localization.

In principle, for situation (2), resolving these coupled effects would require observing the downstream phenotype across a range of controlled upstream values for each variant. For instance, to isolate the effect of protein abundance on surface expression, one would ideally observe surface levels at multiple controlled abundance levels for the same variant, such as through inducible expression systems or titrated mRNA doses. This would allow characterization of the functional relationship between an upstream variable *X*_*v*_ (e.g., abundance) and a downstream phenotype *Y*_*v*_ (e.g., surface expression). However, achieving such finely controlled measurements for thousands of variants in parallel is experimentally infeasible due to technical and biological constraints.

To address this limitation, `Cosmos` introduces position-level aggregation. Rather than estimating causal effects at the level of individual variants *v*, which is under-determined with often only one observation (*X*_*v*_, *Y*_*v*_), we aggregate information across all variants with single mutations that share the same mutated position (*X*_*i*_ and *Y*_*i*_, where *i* indexes protein positions, Fig. 2A). This strategy leverages the biological intuition that amino acid residues with similar structural or functional roles may exhibit coherent downstream behavior once upstream effects are accounted for. For each position *i*, we fit a linear causal model, described in detail in the next section, to estimate the relationship between the upstream and downstream phenotypes. By shifting to the position level, we enable tractable and biologically meaningful inference.

**Fig 2:**
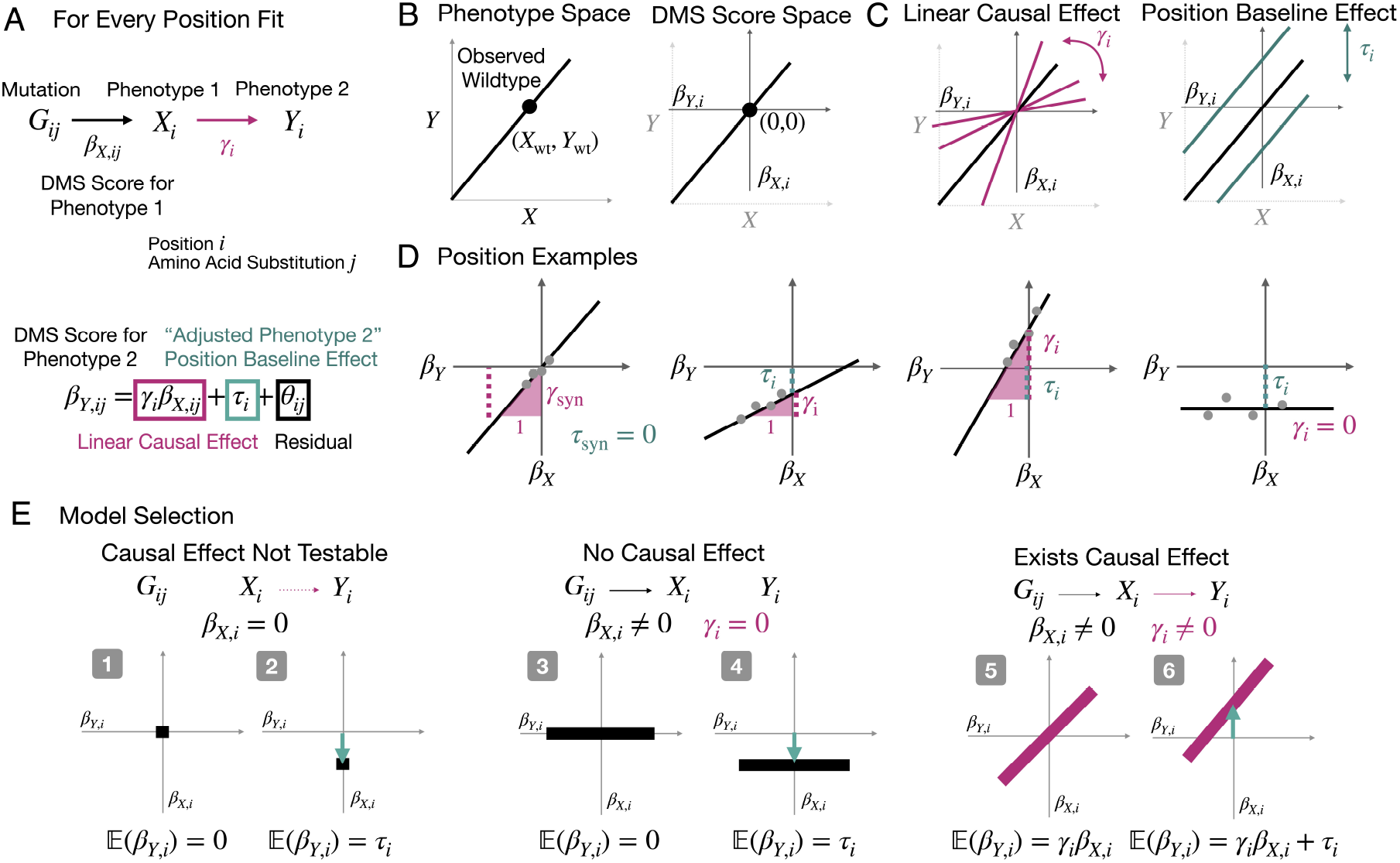
`Cosmos` model specification. A: The linear model underlying `Cosmos` at position-resolution. B: DMS scores are normalized against the wildtype. The DMS score space is effectively a translation of the phenotype space by the observed wildtype phenotype values. C: Illustration of the two terms in the linear model. D: Examples of (*γ, τ*) combinations: *γ >* 0, *τ* = 0; *γ >* 0, *τ <* 0; *γ* ≫ 0, *τ >* 0; *γ* = 0, *τ <* 0. E: Model categories from a causal perspective. The shape of the DMS score scatterplot is determined by the underlying causal structure.

### 2.3 Linear Causal Model on DMS Functional Scores

To estimate the causal relationship between upstream and downstream phenotypes at each protein position, `Cosmos` fits a linear structural equation model that treats the upstream measurement as a causal predictor of the downstream outcome. While biological relationships may be nonlinear, this linear formulation offers a tractable and interpretable approximation that performs robustly across diverse datasets.

Importantly, we do not observe the true phenotypes *X*_*i*_ and *Y*_*i*_ directly. Instead, we work with functional scores, denoted *β*_*X,i*_ and *β*_*Y,i*_, which are estimated from DMS raw measurements through statistical methods and represent transformed versions of the underlying phenotypes. These scores are typically normalized relative to a reference, such as wild-type or synonymous mutations, and are therefore interpreted on a relative rather than absolute scale (Fig. 2B).

The linear structural equation model in `Cosmos` is summarized in Fig. 2A and includes two key parameters. First, the linear causal effect *γ*_*i*_ quantifies the strength of association between the upstream and downstream scores. It can be interpreted as the slope of the regression line, indicating how much the downstream score *β*_*Y,i*_ changes per unit change in the upstream score *β*_*X,i*_ (Fig. 2C). Second, the position-specific baseline effect *τ*_*i*_ represents the expected downstream score when the upstream score is normalized to zero (e.g., matched to wild-type abundance). This parameter serves as an adjusted downstream phenotype baseline, capturing position-specific effects independent of the upstream phenotype (Fig. 2C).

Fig. 2D illustrates theoretical examples of position-level causal fits, showcasing the diversity of functional relationships between phenotypes. (1) At some positions, the regression line passes near the origin (0, 0) with a significant positive causal effect *γ*_*i*_ *>* 0, indicating a proportional relationship between the upstream and downstream phenotypes. (2) Other positions exhibit a reduced slope and a negative baseline effect *τ*_*i*_ *<* 0, indicating that variants at these positions decrease the downstream phenotype even when the upstream variable is held at its reference level, suggesting a downstream-specific loss-of-function mechanism. (3) Some positions show steep positive slopes and elevated intercepts (*γ*_*i*_ ≫ 0 and *τ*_*i*_ *>* 0), consistent with gainof-function variants that enhance the downstream phenotype beyond the upstream influence. (4) Finally, positions with flat regression lines (*γ*_*i*_ = 0) suggest no detectable relationship between the two phenotypes. This may occur when mutations render the protein entirely nonfunctional in the downstream assay, making upstream variation irrelevant.

Another strength of the `Cosmos` framework is its ability to incorporate uncertainty in the functional scores directly into the likelihood function (see Methods). Rather than treating the DMS-derived scores *β*_*X,i*_ and *β*_*Y,i*_ as fixed values, the model accounts for their estimated measurement error, reflecting experimental noise, coverage depth, and replicate variability. This modeling approach allows downstream parameter estimates, including *γ*_*i*_ and *τ*_*i*_, to appropriately reflect the confidence in the underlying data. By integrating uncertainty, `Cosmos` provides a more statistically grounded approach to causal inference, particularly in datasets with uneven data quality.

### 2.4 Bayesian Model Selection in Cosmos Framework

To systematically assign each position to a causal pattern, `Cosmos` uses Bayesian model selection to compare six candidate models that represent distinct combinations of linear causal effect and position baseline shift.

The linear causal effect *γ*_*i*_ represents the effect of upstream variation on the downstream phenotype. At some positions, however, the upstream measurement *β*_*X,ij*_ shows little to no variation across variants, making the causal effect unidentifiable; these positions are considered untestable (Models 1 and 2). For positions with sufficient upstream variation, we distinguish those with no detectable causal effect (*γ*_*i*_ = 0; Models 3 and 4) from those with evidence of a causal relationship (*γ*_*i*_ ≠ 0; Models 5 and 6).

We also jointly test for the position baseline effect *τ*_*i*_, which captures downstream deviations when the upstream variable is at its reference level. For each causal pattern above, the intercept may be absent (*τ*_*i*_ ≠ 0) or present (*τ*_*i*_ = 0), resulting in six total models (Fig. 2E, Table 1). Each model captures a distinct upstream–downstream relationship, ranging from a single point, a flat baseline, to a sloped line with an intercept.

**Table 1:** Summary of the Six (Potentially) Causal Models.

| Model | Upstream effect | Position baseline effect | Causal effect | Scatterplot |
| --- | --- | --- | --- | --- |
| 1 | $\beta_{X_{ij}} = 0$ | $\tau_i = 0$ | Untestable | Point: $(0, 0)$ |
| 2 | $\beta_{X_{ij}} = 0$ | $\tau_i \neq 0$ | Untestable | Point: $(0, \tau_i)$ |
| 3 | $\beta_{X_{ij}} \neq 0$ | $\tau_i = 0$ | $\gamma_i = 0$ | Line: $\beta_{Y_{ij}} = 0$ |
| 4 | $\beta_{X_{ij}} \neq 0$ | $\tau_i \neq 0$ | $\gamma_i = 0$ | Line: $\beta_{Y_{ij}} = \tau_i$ |
| 5 | $\beta_{X_{ij}} \neq 0$ | $\tau_i = 0$ | $\gamma_i \neq 0$ | Sloped line through $(0, 0)$ |
| 6 | $\beta_{X_{ij}} \neq 0$ | $\tau_i \neq 0$ | $\gamma_i \neq 0$ | Sloped line through $(0, \tau_i)$ |

To compare models, we fit each of the six models to the functional score data for a given position and compute their expected log pointwise predictive density (ELPD) [19]. The ELPD quantifies how well a model is expected to generalize to unseen data by integrating over the posterior. Differences in ELPD between models (*Δ*ELPD) are used to select the best-fitting model while accounting for model complexity and uncertainty. This approach allows `Cosmos` to infer, in a data-driven and interpretable way, whether a given residue position supports a downstream-specific effect.

To test the model performance, we simulate data based on a generative model (Methods) to showcase the reliability of `Cosmos` and what factors influence its reliability. In our default test configuration, each position has 20 variants; the noise *σ*_*X,ij*_ and *σ*_*Y,ij*_ in 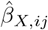 and 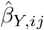 is 15% of the absolute mean of the LOF mixture in the distribution of *β*_*X,ij*_ and *β*_*Y,ij*_ (LOF peak in Gaussian Mixture); and standard error *ω*_*θ*_ of the residual *θ*_*ij*_ is 2.5% of the LOF peak. By this setting, the model selection procedure chooses the true underlying model (from 1 to 6) more than 95% of the time on average. Increasing the magnitude of the noise or decreasing the number of variants per position decreases the success rate, but the performance is still reasonable as long as the noise does not go to more than 50% of the LOF peak (see Supplement Fig. 2).

### 2.5 Position Examples of Kir2.1 Abundance and Surface Expression

We applied `Cosmos` to a deep mutational scanning dataset of the Kir2.1 potassium channel, focusing on two phenotypes: protein abundance and surface expression [10]. A heatmap of both phenotypes of the entire protein sequence is shown in Fig. 3A, revealing distinct spatial patterns and highlighting regions with consistent or divergent functional effects. We highlight six positions across four distinct structural regions of the protein to illustrate the range of causal patterns captured by `Cosmos`.

**Fig 3:**
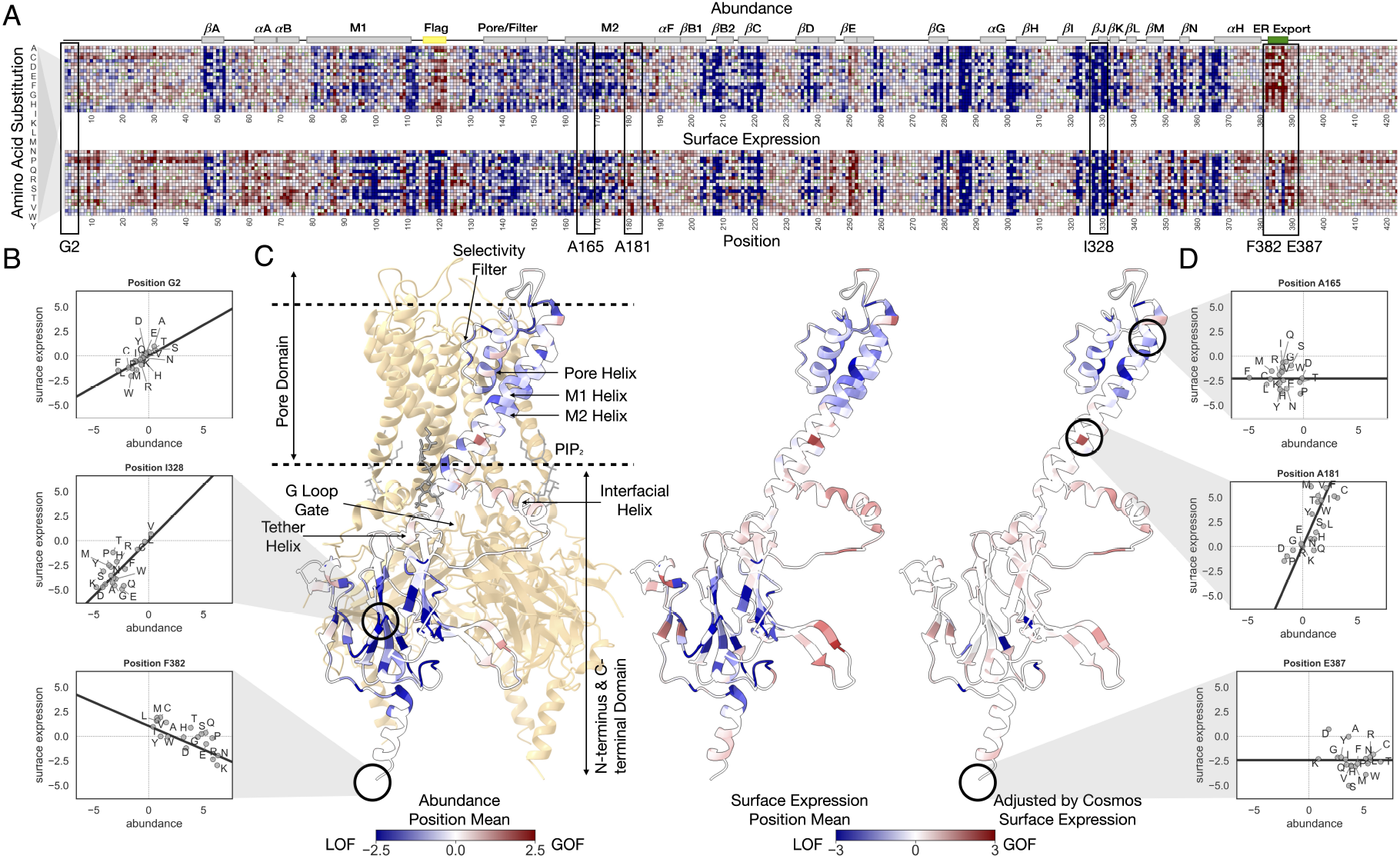
`Cosmos` applied to Kir2.1. A: A heatmap of two of the Kir2.1 phenotypes (protein abundance and surface expression). Black boxes highlight the regions where the following position samples belong. B and D: Scatterplot of two phenotypes of sample positions, showcasing `Cosmos` disentangling various causal patterns at residue-level resolution by selecting different models and distinguishing linear causal effects from baseline effects. The black line denotes the model chosen by `Cosmos` as the best fit. C: Kir2.1 structure visualization, colored by mean abundance, mean surface expression, and the `Cosmos`-adjusted surface expression *τ*_*i*_. Only one monomer of the tetramer is colored in the leftmost panel and shown in the other two. The color schemes of the heatmaps in A and structures in C are the same for the two phenotypes respectively.

- G2, located in the N-terminus, is functionally near-neutral for both phenotypes, and the relationship between abundance and surface expression resembles that of synonymous variants—higher abundance leads to higher surface expression (Model 5, Fig. 3B).
- In the M2 transmembrane helix, A165 fits Model 4: although abundance varies, surface expression remains unchanged, suggesting a decoupling of the two phenotypes (Fig. 3D). By contrast, A181 fits Model 5 but with a steeper slope than G2, indicating that abundance changes are strongly amplified in surface expression (Fig. 3D).
- Position I328, located in the core *β*-sheet of the C-terminal domain (CTD), exhibits loss-of-function in both abundance and surface expression. However, when abundance is elevated to the reference level, surface expression appears near-neutral, where upstream variation fully explains the downstream effect. (Model 5, Fig. 3B).
- Within the ER export signal FCYENE in the C-terminus, both E387 and F382 exhibit elevated abundance. E387 shows no linear causal effect (Model 4, Fig. 3B), as surface expression remains loss-of-function regardless of increased abundance. In contrast, F382 displays a negative causal slope (Model 5, Fig. 3B), indicating that increased abundance is associated with reduced surface expression. This inverse relationship may reflect a trafficking defect, where excess accumulation of protein at this site disrupts proper export.

Together, these examples demonstrate how `Cosmos` disentangles diverse and interpretable causal patterns at residue-level resolution.

### 2.6 Cosmos Reveals Direct Surface Expression Defects at Top Pore Domain in Kir2.1

To distinguish direct surface expression defects from those indirectly caused by reduced abundance, we mapped three position-level metrics onto the Kir2.1 structure (PDB 3SPI, Fig. 3C): mean abundance, mean surface expression, and the adjusted surface expression inferred by `Cosmos` (i.e., the position baseline effect *τ*_*i*_).

In the unadjusted data, both abundance and surface expression show widespread loss-of-function across the core C-terminal *β*-sheets and helices at the top of the pore domain, suggesting that structural perturbations broadly affect protein folding, stability, and trafficking. However, after adjusting for abundance with `Cosmos`, the apparent loss-of-function in the CTD *β*-sheets largely disappears, while strong signal persists in the top pore domain. This pattern indicates that reduced surface expression in the *β*-sheet region is largely an indirect consequence of impaired abundance. In contrast, the persistent signal in the pore helix likely reflects direct disruption of trafficking or membrane localization, highlighting residues in this region as critical for surface expression, independent of upstream abundance effects.

### 2.7 Cosmos Isolates Binding-Specific Effects from Abundance in PSD95-PDZ3 and KRAS

We next applied the `Cosmos` framework to two additional datasets, PSD95-PDZ3 abundance and CRIPT binding [4], and KRAS abundance with RAF1-RBD binding [21], to further evaluate its generalizability beyond the Kir2.1 use case.

In the PSD95-PDZ3 dataset, the scatterplot of abundance versus CRIPT binding displays a structure similar to that of Kir2.1: a well-defined neutral cluster, along with a distinct loss-of-function (LOF) cluster enriched for nonsense mutations (Fig. 4A). To disentangle binding-specific effects from those driven indirectly by abundance loss, we applied `Cosmos`. The resulting model assignments across 84 positions in the domain reveal diverse causal relationships (Fig. 4B). The majority of positions are assigned to Models 5 and 6, indicating clear causal effects of abundance on binding. A few positions are best fit by flat models (Model 3 or 4), particularly at the termini of the domain or near the CRIPT-binding interface (*α*B helix and *β*B sheet), where binding appears decoupled from abundance. Only one position (position 11) is assigned Model 2, lacking sufficient upstream variation to permit causal inference. Mapping the position-level scores—abundance, raw binding, and adjusted binding inferred by `Cosmos` —onto the crystal structure reveals that LOF effects near the CRIPT-binding pocket persist after adjustment, while other regions become largely neutral, highlighting direct binding determinants (Fig. 4C).

**Fig 4:**
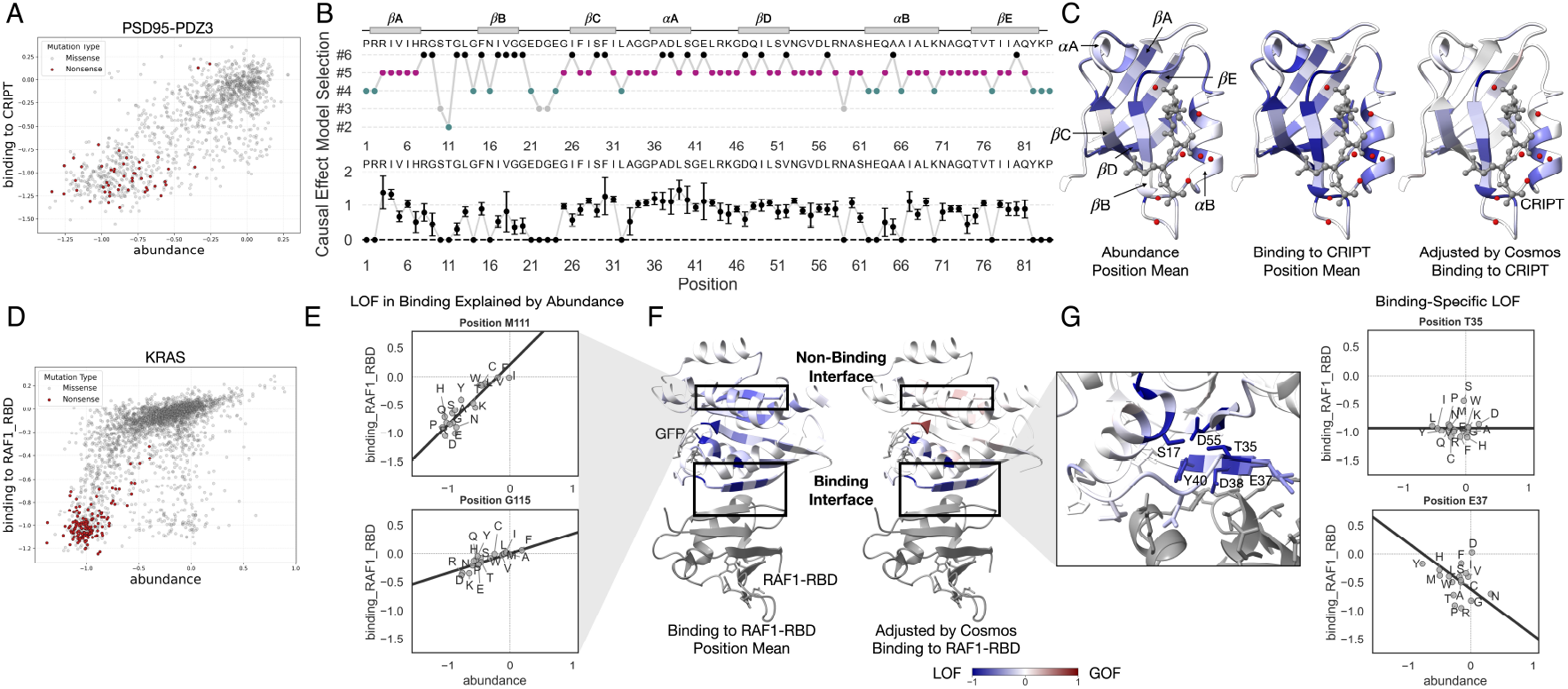
`Cosmos` applied to PSD95-PDZ3 and KRAS. A: Scatterplot of PSD95-PDZ3 abundance and CRIPT binding. Nonsense mutations are colored in red. B: Upper panel: the model label (1-6, see Fig. 2) `Cosmos` assigns to each position of PSD95-PDZ3. Secondary structures are denoted at the top. Lower panel: the causal effect sizes `Cosmos` estimates for each position, with error bars of one standard deviation. C: PSD95-PDZ3 structure visualization (PDB 1be9), colored by mean abundance, mean binding to CRIPT, and `Cosmos`-adjusted binding to CRIPT *τ*_*i*_. Secondary structures are also labeled. The color scale is the same as that in F. D: Scatterplot of KRAS abundance and RAF1-RBD binding. Nonsense mutations are colored in red. E: Scatterplot of example positions in non-binding interface. The black line denotes the model chosen by `Cosmos` as the best fit. F: KRAS structure visualization (PDB 6vjj), colored by mean binding to RAF1-RBD and `Cosmos`-adjusted surface expression *τ*_*i*_. G: A close-up look of KRAS binding interface, highlighting two example positions.

We applied the same procedure to the KRAS dataset, which includes abundance and RAF1-RBD binding measurements. Notably, the KRAS binding screen exhibits strong resolution in separating LOF from neutral variants, with a clearly defined LOF peak (Fig. 4D). However, the dynamic range near the neutral cluster is somewhat compressed, making it difficult to distinguish subtle gain-of-function or mildly deleterious effects.

Mapping the position-level scores onto the KRAS structure reveals a familiar pattern: LOF effects persist at the RAF1-RBD binding interface, while most other regions appear neutral after adjustment by `Cosmos` (Fig. 4F). To illustrate this, we highlight several representative positions. M111 and G115, located outside the interface, exhibit well-fit causal model with a positive slope crossing near the origin, indicating that binding defects are explained by corresponding decreases in abundance (Fig. 4E). In contrast, interface residues T35 and E37 behave differently. E37 lies within a *β*-sheet directly contacting RAF1-RBD [21] (Fig. 4G). T35, though not in direct contact, is spatially proximal. Interestingly, T35, D55, and S17, located on different secondary structural elements, form a physical triad and all exhibit persistent LOF in binding even after adjusting for abundance (Fig. 4G). This pattern suggests that disruption of these interactions may destabilize the adjacent binding interface, leading to loss of binding despite preserved expression.

## 3. Discussion

In this study, we present `Cosmos`, a computational framework for causal analysis of multi-phenotype deep mutational scanning (DMS) data. The framework integrates three key components - position-level aggregation, a linear structural equation model, and Bayesian model selection - to address three central questions in analyzing the multi-phenotype DMS data: (1) Does a causal relationship exist between two phenotypes? This is assessed through model selection among six biologically interpretable scenarios. (2) What is the strength of the causal relationship? This is quantified by the linear causal effect parameter *γ*_*i*_, estimating how changes in the upstream phenotype affect the downstream outcome. (3) What would happen to the downstream phenotype if the upstream phenotype were fixed to a reference value? This is captured by the position-specific baseline parameter *τ*_*i*_, which serves as the adjusted downstream phenotype under a counterfactual upstream condition.

We demonstrated the utility of `Cosmos` through case studies on three proteins: Kir2.1, where we analyzed abundance and surface expression; PSD95-PDZ3, examining abundance and CRIPT binding; and KRAS, exploring abundance and RAF1-RBD binding. Across these examples, `Cosmos` revealed residue-level causal relationships that distinguish direct functional effects from indirect consequences of upstream functional effect, providing biologically interpretable insights that are not accessible through traditional correlationbased approaches.

While `Cosmos` offers a principled approach for inferring causal relationships in multi-phenotype DMS data, several limitations are acknowledged.

First, the method relies on position-level aggregation. Although grouping variants by mutated position increases robustness, it implicitly assumes that variants at the same site share similar causal behavior. This assumption may not always hold: individual substitutions can have distinct structural or biochemical consequences. In such cases, variant-specific modeling, such as in Mochi, would be ideal.

Second, the causal model assumes a linear relationship between upstream and downstream phenotypes. However, biological systems may be nonlinear due to assay saturation, threshold effects, or intrinsic nonlinear dynamics, such as the preset biophysical nonlinear model used by Mochi. When the overall relationship deviates substantially from linearity, methods that explicitly allow for learnable nonlinear trends may be more appropriate and could be a future direction for extension.

Third, `Cosmos` performs best when the dataset spans a wide dynamic range across phenotypes—capturing loss-of-function (LOF), neutral, and potentially gain-of-function (GOF) variants. In datasets with limited dynamic range, where most variants cluster tightly around a baseline, simple outlier detection may be more appropriate than causal modeling.

Furthermore, `Cosmos` is constrained to the analysis of two phenotypes (downstream and upstream), but (1) there could be more than two phenotypes measured, or (2) the relationships are not sequential but parallel (different binding ligands in a binding screen) or multi-conditioned (different concentrations of ligand as a response curve). Methods specific to other phenotype relationships require further development.

While not solving all the problems, `Cosmos` establishes a principled and interpretable framework for residue-level causal inference in multi-phenotype deep mutational scanning, offering a valuable tool for dissecting the molecular basis of protein function across diverse biological contexts.

## 4 Methods

### 4.1 Notation and Linear Structural Equation Model

*X* (upstream) and *Y* (downstream) are the two phenotypes of interest. Each single-mutation variant *v* is identified by its position *i* and amino acid substitution *j*. The “true” (unobserved) functional scores of the two phenotypes are denoted *β*_*X,ij*_ and *β*_*Y,ij*_. The mean estimates and standard errors of such effects from the statistical analysis fo the raw DMS measurements are 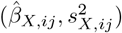 and 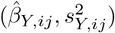 respectively.

Assuming linear causal effects, the true effect of *Y, β*_*Y,ij*_, can be decomposed as

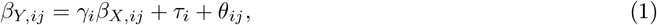

with each term standing for: *γ*_*i*_ as linear causal effect, *τ*_*i*_ as position baseline effect, and *θ*_*ij*_ as remaining residual.

### 4.2 Likelihood and Prior of the Six Models

The joint distribution of true functional score *β*_*X,ij*_ and the residual *θ*_*ij*_ is modeled as a Gaussian mixture. We use ∑_*k*_ *π*_*k*_N_2_ to denote a bi-variate Gaussian random variable with *k* mixtures (instead of a weighted sum of Gaussian random variables):

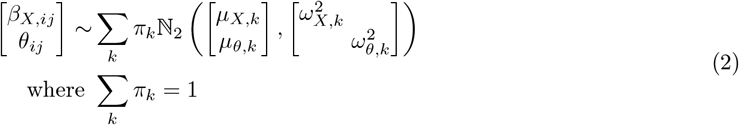

Combining Equations 1 and 2, we obtain the joint distribution of the true functional scores of both phenotypes:

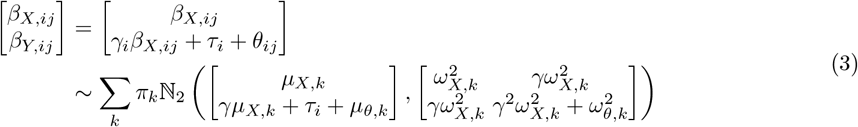

The mean estimates from the statistical analysis 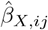 and 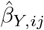 are noisy measurements of the true functional scores:

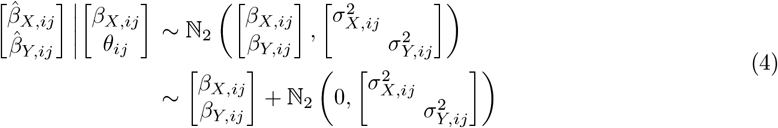

The unconditional joint distribution of the mean estimates is thus:

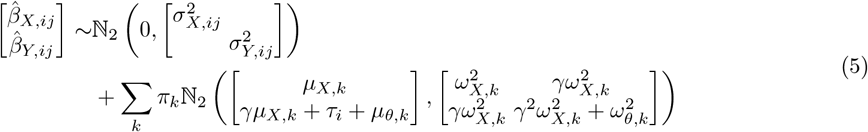

Since the latent variables 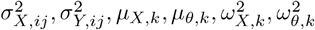 are unobserved, we replace them with estimated quantities:

- 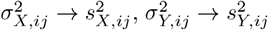: Each observation’s noise magnitude is replaced with standard error of effect estimates.
- 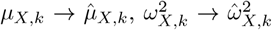: Estimated by fitting effect estimates of 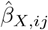 to a Gaussian mixture, with compensation for the noise term *σ*^2^ (see below).
- *µ*_*θ,k*_ → 0: Residual has zero mean by construction.
- 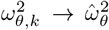: Replace with the variance of the effect estimate of 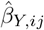, multiplied by a constant to account for the contribution of other variance terms. This is a heuristic treatment as the variance of 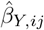 has an unknown component *γ*.

We further simplify the problem by assuming that *θ* follows a single-component Gaussian mixture (the Gaussian distribution itself), resulting in *β*_*X,ij*_ and *θ* being independently distributed (but not *β*_*X,ij*_ and *β*_*Y,ij*_):

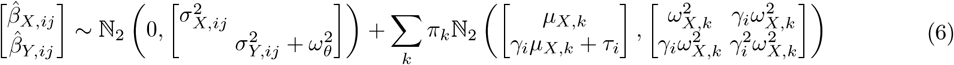

Estimating *µ*_*X,k*_ and 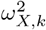 rigorously requires fitting the Gaussian mixture on 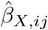 with heteroskedastic but known noise 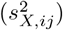, which could involve a customized Expectation-Maximization algorithm or even fitting a Bayesian hierarchical model. We opt for a simpler and faster approach, by directly fitting a Gaussian mixture on 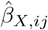, getting 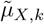 and 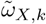, and then adjusting the variance term by the geometric mean of all *s*^2^:

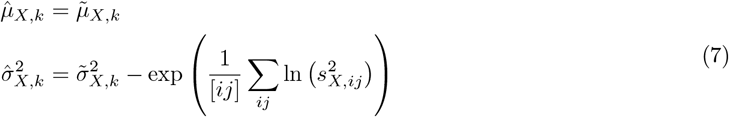

Fitting the Gaussian mixture on 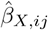 also identifies which component each variant belongs. Denote the estimated mapping by 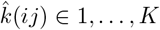. Note that *γ*_*i*_ and *τ*_*i*_ do not determine the distribution of 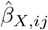 and consequently our categorization of variants.

We have thus obtained the (empirical) likelihood function for each observed effect estimate:

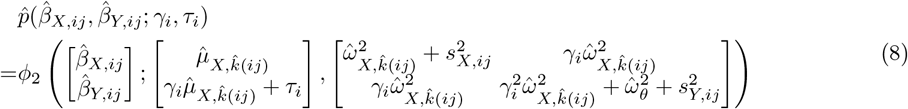

where *ϕ*_2_(·; *µ, Σ*) is the probability density function of a bivariate Gaussian random variable with mean vector *µ* and covariance matrix *Σ*.

*τ*_*i*_ and *γ*_*i*_ are given Gaussian but very flat prior independently.

The prior and the likelihood function above corresponds to Model 6, the full model. Models 3 to 5 share the same likelihood function with restrictive prior on certain parameters: Model 3 (*γ* = *τ* = 0); Model 4 (*γ* = 0); Model 5 (*τ* = 0).

Models 1 and 2 assume that the upstream effect and the amino-acid substitution effect are both zero (*β*_*X,ij*_ = *θ*_*ij*_ = 0). As a result, the causal effect *γ*_*i*_ is also absent from the likelihood function of effect estimates:

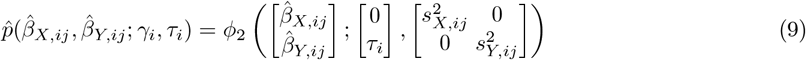

Model 1 further restricts that *τ*_*i*_ = 0.

In addition, if the standard error 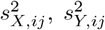 are not available, we will drop the noise term 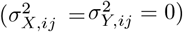 and fit Models 3-6 only.

### 4.3 Adaptive Grid for Posterior Approximation

With the prior and the likelihood function, we can approximate the posterior for each model using an adaptive grid. We implemented an adaptive grid algorithm, adapted from `CAUSE`[11], to approximate the posterior over any reasonable number of dimensions. It automatically expands the sampling area before adaptively refining certain hypercubes in the grid, each controlled by a user-defined tolerance.

We fit all six models on each of the positions (*i*). To show how our adaptive grid works, take Model 6 and position *i* = 0 as an example. The parameters of interest are *γ*_0_ and *τ*_0_, the causal effect and the position baseline effect of Position 0. Running a linear regression of *β*_*X*,0*j*_ against *β*_*Y*,0*j*_, we get the estimated slope 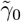 and intercept 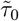, from which we set the initial range of each parameter 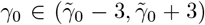 and 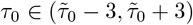. Each axis has an interval size of 0.05.

The prior over each interval on each axis is evaluated to obtain the prior on each hypercube in the grid. The likelihood function is evaluated on the midpoint of each hypercube and assumed to be uniform over the entire hypercube. The log posterior function is estimated by:

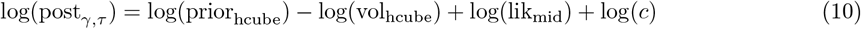

where each term stands for:

- post_*β*_: The posterior function evaluated at (*γ, τ*),
- prior_hcube_: The prior over the entire hypercube to which (*γ, τ*) belongs,
- vol_hcube_: The volume of the hypercube,
- lik_mid_: The likelihood function evaluated at the midpoint of the hypercube,
- *c*: Normalizing constant.

If the marginal posterior probability of an axis’s lowest or highest interval is higher than a user-defined threshold (`ADAPT_BOUNDARY_THRES`), the adaptive grid will be expanded along the axis in that direction. The posterior function is re-approximated before continuing with the next direction or axis. We cycle through the two directions of all axes until none of the axes require expansion anymore (Supp. Fig. 1.1A).

**Supplementary Figure 1.1:**
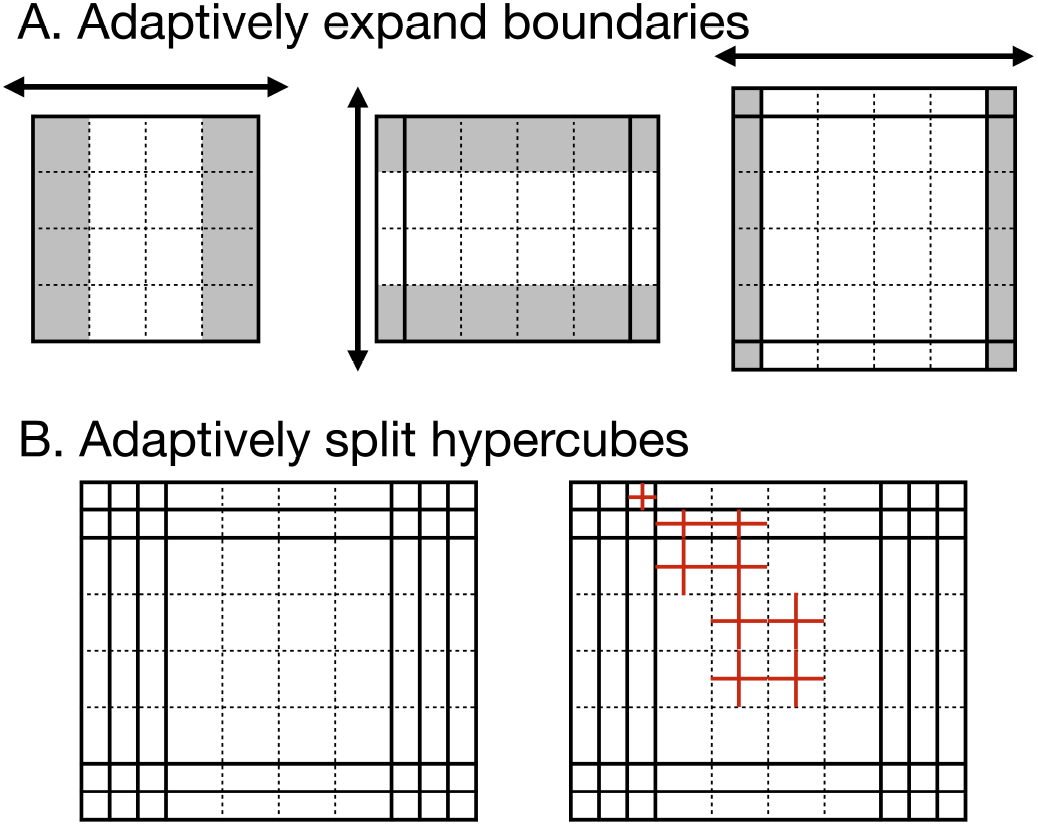
Adaptive grid expansion and bisection. A: Expand the boundaries if marginal posterior on either side of an axis (colored in gray) exceeds a threshold, cycling through all axes multiple times. B: After the boundaries are adaptively expanded, hypercubes are split by bifurcating each edge, until none of the hypercubes has an estimated posterior exceeding a threshold.

After grid expansion, we scan for hypercubes with log posterior (log(post_hcube_) = log(post_*β*_)+log(vol_hcube_)) exceeding the grid splitting threshold (ADAPT_SPLIT_THRES). All such hypercubes are divided into 2^*n*^ (each edge is bifurcated). The grid will be repeatedly scanned and refined until no more such hypercubes are found (Supp. Fig. 1.1B).

The adaptively expanded and refined grid can readily evaluate the posterior for any given (*γ, τ*) and generate draws of (*γ, τ*) according to the posterior.

### 4.4 Bayesian Model Selection

After we fit all the models for a position, they are compared using expected log pointwise predictive density (ELPD) [19]. We first generate a sample for each model (1000 random draws from the grid described above). Given a log-likelihood function, we can estimate the ELPD for any sample of parameters. The models are then compared and ranked by their samples’ ELPD.

Our estimation of ELPD uses the Pareto smoothed importance sampling leave-one-out cross-validation, “PSIS-LOO-CV”, implemented by the `ArviZ` package [8]. An issue of concern is that PSIS-LOO-CV could fail to produce a reliable estimation of ELPD. Our package relays the warning produced by `ArviZ`. In our analysis of proteins such as Kir2.1, the warnings only occur sporadically.

It is also possible that we estimate the ELPD of a model fit on one position with the log-likelihood function generated by another position, to evaluate the cross-validity of models. While it is not covered in this paper, we have implemented the functionality in our package.

### 4.5 Simulated Data Generation

We devised a generative model to simulate data for evaluating the accuracy and efficacy of `Cosmos`. In our default test configuration, we simulated 50 positions, each with 20 variants. Position-level and variant-level information were generated sequentially.

First, each position *i* was assigned a model index, either `null` (corresponding to Models 1 and 2) or `mixed` (corresponding to Models 3 through 6). Next, we sampled the position-level parameters *γ*_*i*_ and *τ*_*i*_ from a two-component Gaussian mixture model. One of the mixture components was degenerate and fixed at 0. Specifically, Models 2, 4, and 6 correspond to a degenerate mixture in *τ*_*i*_, and Models 1, 2, 3, and −4 correspond to a degenerate mixture in *γ*_*i*_. The significant mixture component for *γ*_*i*_ was defined by a normal distribution with mean 1 and standard deviation (sd) 0.3, with a mixture proportion of 0.4. For *τ*_*i*_, the significant mixture had a mean of 4 with sd 2 and a mixture proportion of 0.6. Together, the model index and the sampled parameters *γ*_*i*_ and *τ*_*i*_ define the “ground truth” model label for each position.

For the variant-level information, we draw each *β*_*x,ij*_ from a two-component Gaussian mixture model, representing a neutral peak (mean 0, sd 0.5, proportion 0.6) and a LOF peak (mean −2, sd 1). Using the position-level parameters *γ*_*i*_ and *τ*_*i*_ and incorporating an additional noise term *θ*_*ij*_ from a normal distribution with sd *ω*_*θ,ij*_ = 0.05, we computed the “true” downstream functional scores *β*_*y,ij*_ using Equation 1. Finally, we add experimental noise to simulate observed effects. Specifically, we apply additive Gaussian noise with sd *σ*_*X,ij*_ = 0.3 and *σ*_*Y,ij*_ = 0.4 for the upstream and downstream phenotypes, respectively, yielding the observed scores 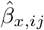 and 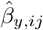.

Starting from this initial configuration, we repeatedly double one or all of the observation noise magnitudes and shrink the number of variants per position. As expected, the predictive power of `Cosmos` gradually decays as observation noises are amplified or the sample size is decreased (Supp. Fig. 1.2).

**Supplementary Figure 1.2:**
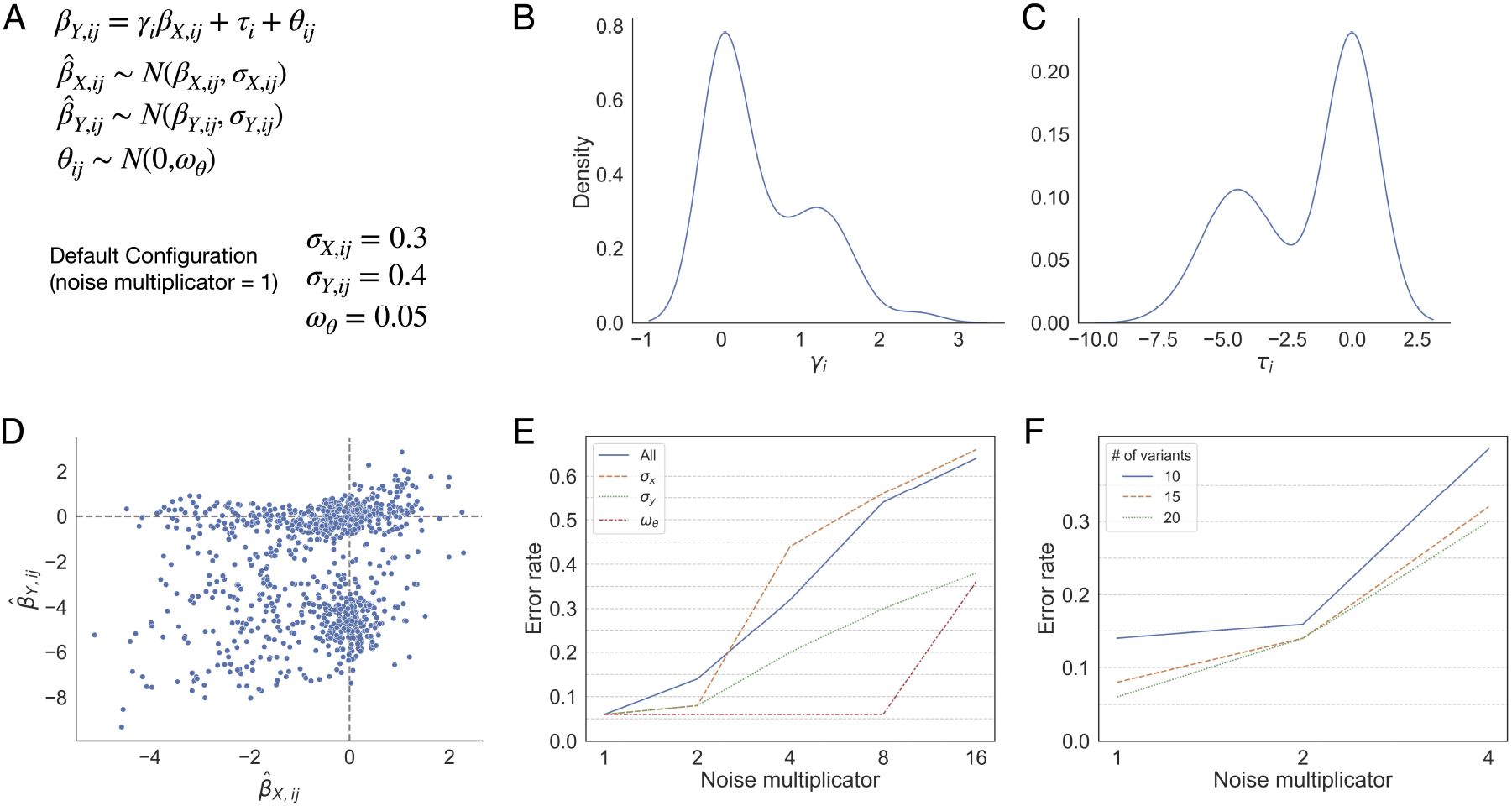
A: Generative model and default hyperparameter configuration. B, C: The distribution of *γ* and *τ* under the default configuration. D: The joint distribution of 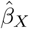 and 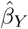 under the default configuration. E: Model selection error rate if the specified noise is multiplied by the multiplicand. F: Model selection error rate by the number of variants per position (coverage).

## 5 Data Availability

`Cosmos` is implemented as a Python package and is distributed on GitHub (https://github.com/pimentellab/cosmos), under the MIT open-source license. The package is also available on PyPI (https://pypi.org/project/cosmos-dms/1.0.1/).

Scripts and pre-processed public datasets used to perform data analysis and generate figures for the paper are uploaded on GitHub (https://github.com/roserao/cosmos-paper-script.git).

The deep mutational scanning datasets we used are as follows: Kir2.1 [10], PSD95-PDZ3 [4], and KRAS [21]. All data are available as supplementary files to their respective publications.

## Acknowledgments

HP is supported by the HHMI Hanna H. Gray Fellowship. WCM is supported by the Hypothesis Fund award and the Shurl and Kay Curci Foundation research grant. MH is supported by the NIH National Institute of General Medical Sciences 5T32GM139786.

## Author Contributions

JR, MH, WCM, and HP jointly conceived the project. JR and HP developed the statistical model. JR developed the simulation framework. JR and MW wrote the software and its support. JR performed the data analysis on protein examples. JR wrote the manuscript with input from MW, MH, WCM, and HP. All authors read and approved the final manuscript.

